# Probing the mechanism of peptidoglycan amidase activation by FtsEX-EnvC

**DOI:** 10.1101/2025.04.02.646785

**Authors:** Jonathan Cook, Allister Crow

## Abstract

The FtsEX-EnvC-AmiA/B system is a key component of the *E. coli* cell division machinery that directs breakage of the peptidoglycan layer during separation of daughter cells. Structural and mechanistic studies have shown that ATP binding by FtsEX in the cytoplasm drives periplasmic conformational changes in EnvC that lead to the binding and activation of peptidoglycan amidases such as AmiA and AmiB. The FtsEX-EnvC amidase system is highly regulated to prevent cell lysis with at least two separate layers of autoinhibition that must be relieved to initiate peptidoglycan hydrolysis during division. Here we test the FtsEX-EnvC amidase activation mechanism through site-directed mutagenesis. We identify mutations that disrupt the autoinhibition mechanism of FtsEX-EnvC and an N-terminal deletion variant that prevents activation. Finally, we develop a cysteine locking residue pair that stabilises the complex in its amidase activating conformation. The reported EnvC variants greatly enhance our understanding of the FtsEX-EnvC autoinhibition mechanism and the conformational changes underpinning amidase activation. Our observations are consistent with the proposed mechanism of amidase activation by large-scale conformational changes in FtsEX-EnvC allowing recruitment and activation of peptidoglycan amidases.

**Importance:** In *E. coli*, the FtsEX-EnvC system regulates two of the three division-associated amidases that break the peptidoglycan layer during bacterial division. Structural and mechanistic studies have revealed a detailed molecular mechanism for amidase activation in which an ABC transporter and its periplasmic partner reversibly activate periplasmic amidases under direction of the cytoplasmic cell division machinery. This paper explores structural features of EnvC that underpin autoinhibition and the activation mechanism. The FtsEX-EnvC system serves as a powerful example of a Type VII ABC transporter that uses transmembrane conformational change to drive work in the periplasmic space.

## Introduction

The peptidoglycan layer is a core component of the bacterial cell envelope that provides both shape and rigidity as well as a point of attachment for the outer membrane (1–5). During cell division there is a need to break the peptidoglycan layer so that newly formed daughter cells can be separated from one another (6). Breaks in the peptidoglycan layer are also required to expose new connection points from which additional peptidoglycan can be added during bacterial elongation (3). In *E. coli*, two division-associated peptidoglycan amidases (AmiA and AmiB) are each located in the periplasm and are regulated by a Type VII ABC transporter complex: FtsEX-EnvC (7–9). FtsEX-EnvC is recruited to the Z-ring early in division (10,11) and regulates the activity of AmiA and AmiB through transmembrane conformational changes driven by FtsEX (8,9,12,13).

*E. coli* strains lacking FtsE, FtsX or EnvC typically form long chains of cells that are connected by the unbroken peptidoglycan layer (13). These strains are susceptible to antibiotics and detergents that would not normally cross the outer membrane barrier and have reduced viability on low salt media indicating cell envelope defects (8,11,14–16). Such phenotypes are best understood as the result peptidoglycan defects arising from the absence of proper amidase activation. However, amidase variants have also been reported with promiscuous activity that produce partial cell lysis (9,17,18). Both inhibition of amidase activation or promiscuous activation of amidases have therefore been seen as potential routes for developing useful antimicrobials targeting cell envelope integrity and bacterial cell division.

A molecular mechanism for the *E. coli* FtsEX-EnvC-AmiA/B system has been developed and is depicted in **Fig. 1a**. In their resting states, periplasmic amidases such as AmiA and AmiB are unable to bind and hydrolyse peptidoglycan because an internal autoinhibitory helix blocks access to their respective active sites (9,18,19). Activation of AmiA or AmiB requires a conformational change in the amidase that displaces the amidase autoinhibitory helix and allows rearrangement of residues around the active site zinc ion (9). The conformational changes in AmiA or AmiB are induced by the binding of a cognate murein hydrolase activator (EnvC) (8,9,20–22). A crystal structure of a periplasmic amidase bound to the EnvC LytM domain has shown how the amidase interacts with a specific binding groove in the EnvC LytM domain (9). Binding of the amidase’s ‘interaction helix’ causes displacement of the amidase’s ‘blocking’ helix exposing the active site to the peptidoglycan layer (9). EnvC is also subject to its own autoinhibition mechanism (8) and is bound to the periplasmic domains of a Type VII ABC transporter (FtsEX) (8,12,13). In its resting state, the EnvC amidase binding site (located within its C-terminal LytM domain) is blocked by a long helix termed the restraining arm, that must be displaced to allow the binding of amidases (8). As depicted in the figure, ATP binding in the cytoplasm drives a transmembrane conformational change that propagates through both FtsEX and the EnvC coiled coil domain displacing its restraining arm and facilitating the binding of amidases (**Fig. 1a**). ATP hydrolysis returns the system to the inactive resting state after a period of activation. The FtsEX-EnvC-AmiA/B system represents a remarkable example of conformational change with multiple levels of autoinhibition that protect the cell from improper peptidoglycan hydrolase activity.

**Figure 1.**
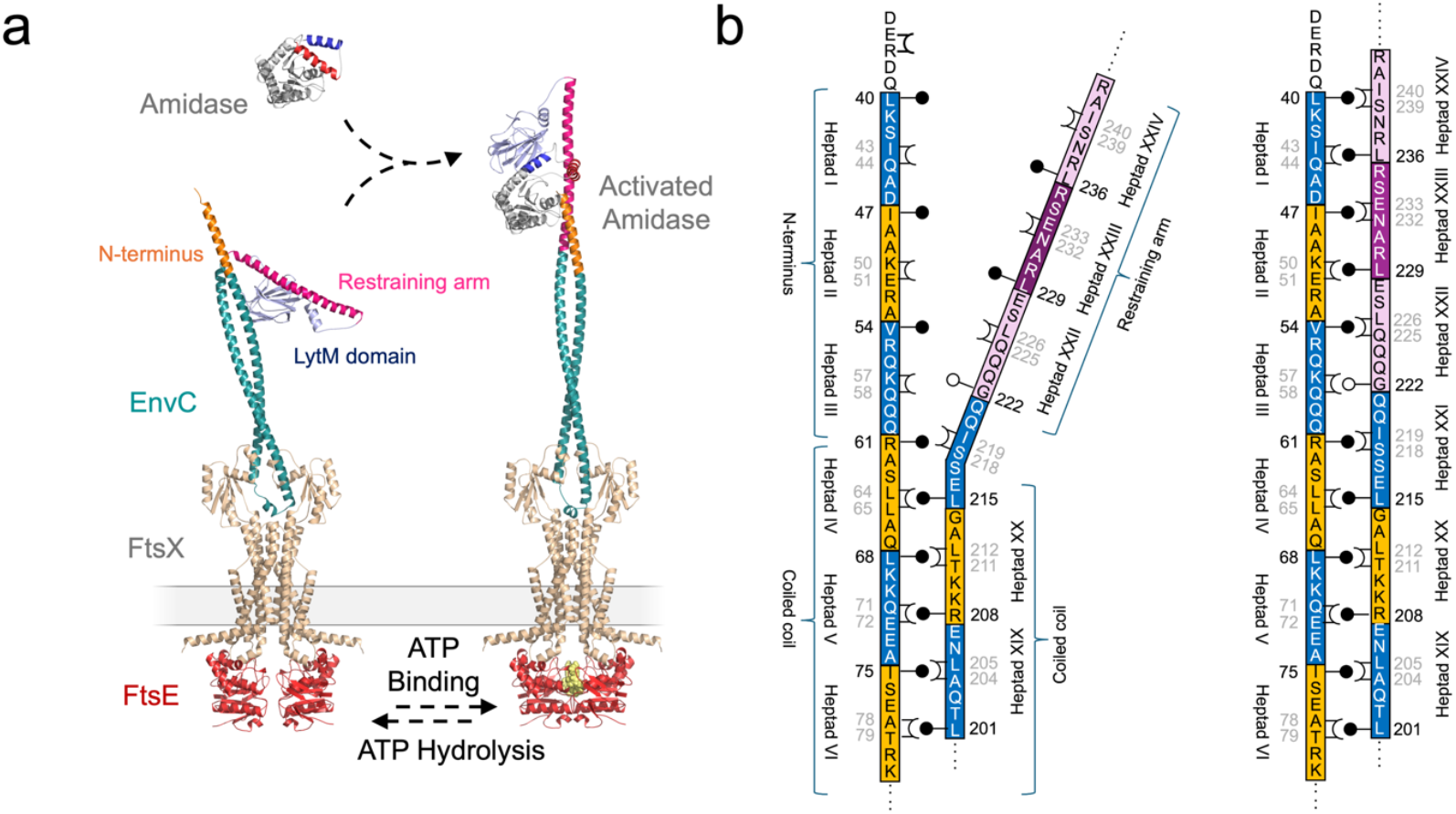
Mechanism of amidase activation by *E. coli* FtsEX-EnvC. (a) ATP binding to FtsE in the cytoplasm causes a transmembrane conformational change to propagate through the FtsEX-EnvC complex leading to the binding and activation of a peptidoglycan amidase. ATP hydrolysis resets the system causing the release (and deactivation) of the amidase as the FtsEX-EnvC complex returns to its inactive resting state. Models shown are hypothetical (ie modelled rather than experimentally determined) but based on structures that have been described previously. (b) Diagram showing the proposed knobs-into-holes packing between the EnvC N-terminus and the restraining arm. Knobs indicated as black pins, and holes are indicated with crowns. G222 is indicated by a white-filled pin to emphasise the absence of a sidechain in this knob position.

Activation of EnvC relies on the elongation of its coiled coil domain and involves multiple heptad repeats in the protein N-terminus and the restraining arm that are predicted to be paired up in the active conformation but parted in the resting state (8) (**Fig. 1b**). Formation of the extended coiled coil during activation is supported by the identification of co-evolving residue pairs between the restraining arm and N-terminus and identification of residues that would support classical knobs-into-holes packing (8,23) (**Fig. 1b**). The conformational change mechanism is broadly supported by observations from various partial structures of the FtsEX-EnvC-AmiA/B complex (8,9,18,22), cryoEM data (24–28) and hypothetical models (8,9), however, the importance of the reversible formation of an extended coiled coil remains to be tested.

Here we examine the importance of three structural features within EnvC: (1) the restraining arm responsible for autoinhibition; (2) a potential molecular hinge between the restraining arm and the coiled coil domain; and (3) the N-terminus of EnvC which supports the formation of the ‘active’ amidase-recruiting conformation through the formation of an extended coiled-coil. Using a combination of *in vitro* and *in vivo* work, we identify mutations that disrupt the autoinhibition mechanism and test the importance of the proposed reversable knobs-into-holes packing that underpins proposed conformational changes in EnvC. These data enhance our understanding of the EnvC autoinhibition mechanism and the structural features underpinning amidase activation by the FtsEX-EnvC-AmiA/B system.

## Results

### Mutations in the restraining arm disrupt the autoinhibition mechanism of EnvC

The EnvC restraining arm is a key structural element that prevents the binding and activation of peptidoglycan amidases (8). In its resting state, the restraining arm of EnvC blocks access to the amidase binding groove located in the LytM domain preventing amidase activation. However, in the activated conformation, the restraining arm is displaced allowing amidases to bind (8,9). A pair of hydrophobic residues (Leu236 and Ile240) each appear to be important for maintaining the restraining arm in the autoinhibited state as they lock inside the hydrophobic groove that would otherwise accommodate the amidase (**Fig. 2a**).

**Figure 2.**
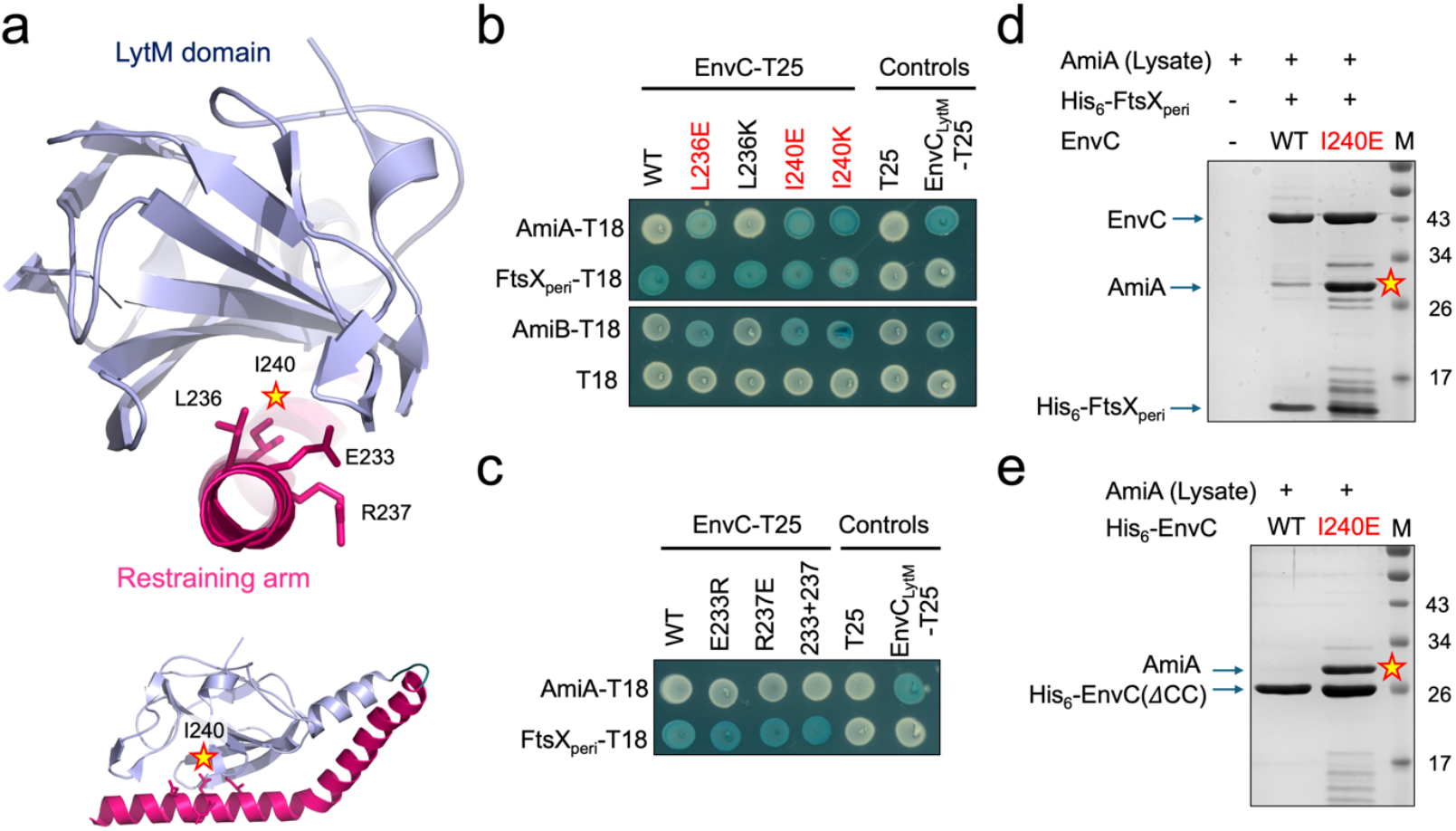
Mutations in the EnvC restraining arm overcome autoinhibition and promote amidase binding. (a) Structure of EnvC with focus on the restraining arm. Stars indicate the locations of the autoinhibition-breaking mutations identified herein. (b) Bacterial 2-hybrid assay testing the interaction between EnvC variants and the two division-associated periplasmic amidases (AmiA and AmiB). The interaction with the FtsX periplasmic domain forms an additional control to show that all full length EnvC proteins are expressed and folded. Red lettering indicates EnvC mutations that overcome autoinhibition and allow promiscuous amidase binding. (c) Bacterial 2-hybrid assay probing the interaction between EnvC variants carrying restraining arm mutations that do not break autoinhibition. (d) Comparative co-purification of AmiA with a soluble complex composed of EnvC and the FtsX periplasmic domain. Only the FtsX periplasmic domain is His-tagged. A star indicates a dense AmiA band arising from copurification with I240E EnvC. (e) Comparative co-purification of AmiA with truncated forms of EnvC that lack the N-terminal coiled coil domain but retain both the LytM domain and restraining arm. A star indicates a dense AmiA band that only co-purifies with the I240E variant.

We predicted that substituting hydrophobic residues for charged residues might release the restraining arm and overcome the EnvC autoinhibition mechanism. To test this, we used a bacterial 2-hybrid experiment monitoring the interaction between EnvC and its cognate amidases, AmiA and AmiB and made either lysine or glutamate mutations in Leu236 and Ile240. In line with previous observations (8), neither AmiA nor AmiB interact with the wild type EnvC (consistent with its autoinhibition by the restraining arm) while the isolated EnvC LytM domain, for which the restraining arm is absent, interacts with both (**Fig. 2b**). In keeping with our predictions, we found that L236E, I240E and I240K EnvC variants all interact with both AmiA and AmiB (**Fig. 2b**).

Having established that the hydrophobic residues at positions 236 and 240 are needed for autoinhibition, we next considered two nearby charged residues (Glu233 and Arg237) that are located on the restraining arm but face away from the LytM domain. These residues form useful counter-predictions as they are not expected to be important for maintaining autoinhibition but are positioned close to Leu236 and Ile240. Consistent with their structural context, Glu233Arg or Arg237Glu substitutions do not disrupt autoinhibition and these EnvC variants remain unable to bind AmiA or AmiB in the bacterial 2-hybrid assay – even though both variants interact with the periplasmic domain of FtsX suggesting they are competently expressed and folded (**Fig. 2c**). Taken together, these data show that L236E, I240E and I240K mutations on the restraining arm all disrupt the EnvC autoinhibition mechanism allowing binding of peptidoglycan amidases.

### Characterisation of the I240E autoinhibition-breaking mutation

Having identified mutations that break the EnvC autoinhibition mechanism in the bacterial 2-hybrid assay, we selected our most effective mutation, I240E, for further characterisation *in vitro*. We have previously shown that EnvC can be co-purified with the truncated periplasmic domains of FtsX and solved a crystal structure of this complex (8). However, the truncated FtsX-EnvC complex does not readily bind to either AmiA or AmiB due to the EnvC autoinhibition mechanism. We therefore tested whether introduction of the I240E mutation disrupts autoinhibition and allows co-purification of peptidoglycan amidases with the truncated FtsX-EnvC complex (**Fig. 2d**). We first mixed cell lysates containing the His-tagged soluble FtsX-EnvC complex and AmiA (without a His-tag) and immobilised the resultant complexes on Ni-IMAC resin. After washing, the remaining proteins were eluted from the resin and analysed by SDS-PAGE to identify potential amidase-activator complexes. As a control, we also performed the experiment using the AmiA lysate without any FtsX-EnvC present. As expected, untagged AmiA does not bind the empty IMAC resin, and has only a weak interaction with the wild type FtsX-EnvC complex. However, introduction of the I240E mutation in the truncated FtsX-EnvC complex produced a much stronger interaction with AmiA consistent with disruption of autoinhibition and unimpeded binding of the amidase (**Fig. 2d**).

To further probe the I240E mutation, we explored amidase binding to a minimal protein construct that is composed of only the EnvC LytM domain and restraining arm (**Fig. 2e**). The EnvC(ΔCC) construct lacks the N-terminal coiled coil domain but is expected to remain autoinhibited due to the presence of the restraining arm. We found no evidence of AmiA binding to the wild type EnvC(ΔCC) construct, but strong binding to its I240E variant (**Fig. 2e**). These results demonstrate the importance of Ile240 in maintaining EnvC in its inactive conformation and the ability of the I204E mutation to overcome autoinhibition.

### The I240E mutation causes growth defects when expressed in the periplasm

Having shown that the I240E mutation in the restraining arm causes EnvC to readily bind amidases in both the bacterial 2-hybrid and co-purification assays, we next tested whether this mutation has a disruptive effect *in vivo*. Our prediction was that the I240E variant should promiscuously bind and activate amidases causing outer membrane defects and impairing bacterial growth. To test, we expressed the I240E EnvC variant in the periplasm of *E. coli* and monitored bacterial growth in LB broth. Periplasmic expression of the wild type EnvC protein had no effect on bacterial growth compared to the negative control, however expression of the I240E EnvC variant significantly impaired growth (**Fig. 3a** *left*). When challenged with broth containing SDS detergent, growth was further reduced for the strain expressing the EnvC I240E variant consistent with outer membrane defects (**Fig. 3a** *right*). We also repeated these experiments using an EnvC knockout strain (**Fig. 3b**). Strains lacking the chromosomal copy of *envc* exhibit modest growth impairment and severe detergent sensitivity - with both phenotypes rescuable by expression of the wild type EnvC from a plasmid (**Fig. 3b**). However, when the I240E variant was introduced into the *envc* knockout strain, the growth defect was intensified (**Fig. 3b** *left*) and the cells remained detergent sensitive (**Fig. 3b** *right*). Taken together, these data are consistent with our characterisation of I240E as a mutation that disrupts the EnvC autoinhibition mechanism and facilitates promiscuous binding and activation of peptidoglycan amidases in the periplasm.

**Figure 3.**
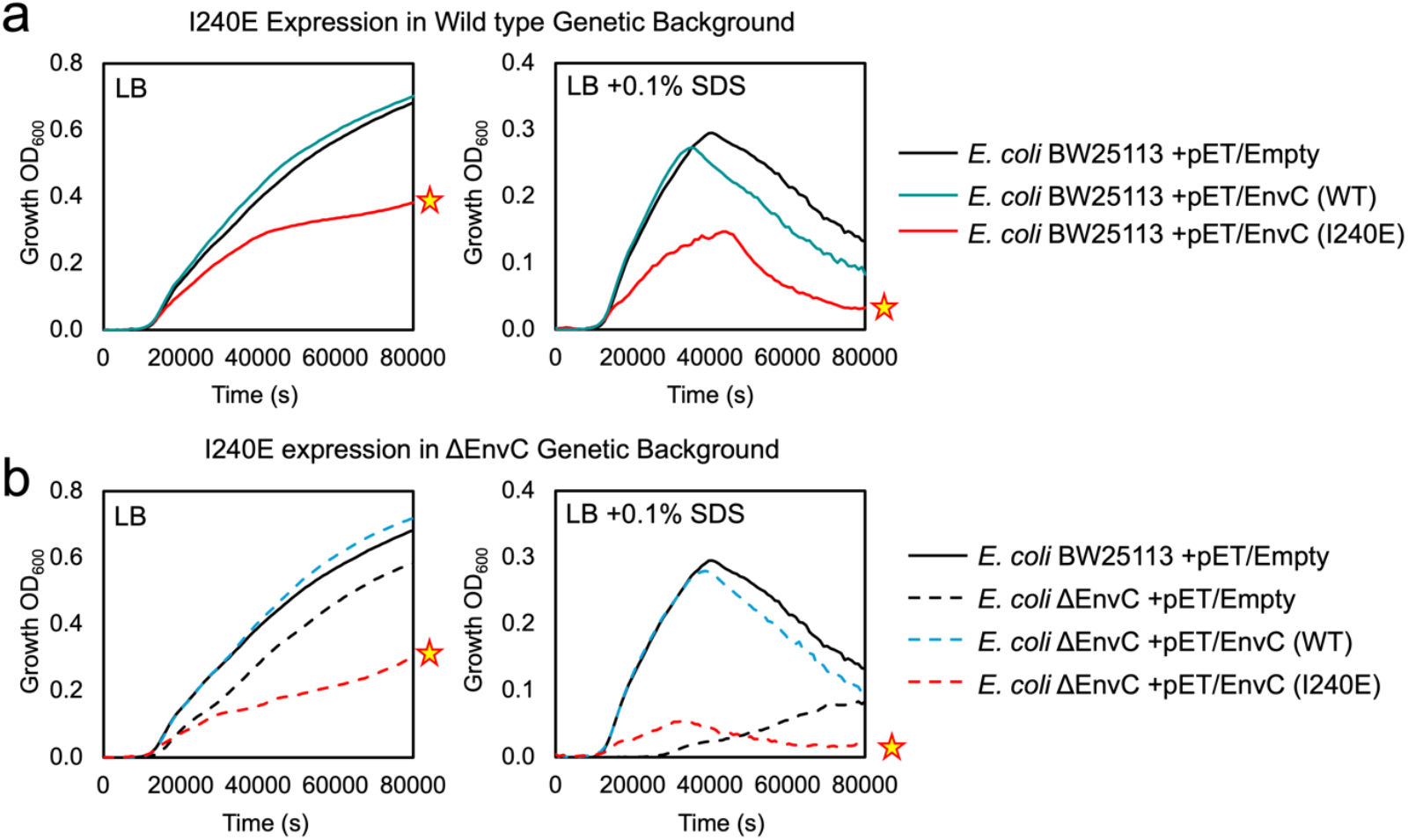
The EnvC I240E variant shows growth defects and SDS sensitivity. (a) Growth curves for *E. coli* BW25113 in LB broth (left) and LB broth supplemented with SDS (right). Strains each harbour a different pET22-based plasmid expressing the indicated EnvC variant. A strain carrying the empty vector is used as a control. A star indicates the impaired growth curve due to expression of the EnvC I240E variant. (b) A similar set of growth curves using an isogenic strain that lacks the chromosomal *envc* gene. All growth curves are the mean of multiple repeats.

### Mutations targeting a proposed molecular hinge in EnvC

Structures of EnvC in its autoinhibited conformation (8) suggests the presence of a ‘hinge’ region connecting the N-terminal coiled coil domain with the restraining arm (**Fig. 4a**). During activation, the hinge is expected to transition between amidase-activating and autoinhibited conformations. In the autoinhibited state, the hinge is bent as per the crystal structure of EnvC, but in the activated state, it is predicted that these residues adopt a continuous helical conformation that links the coiled coil with the restraining arm. Straightening of the hinge thus allows knobs-into-holes packing between heptads in the restraining arm and the N-terminus. We identified a glycine residue (G222) that might provide intrinsic flexibility to the hinge (**Fig. 4a,b**). Gly222 is present among many enterobacterial EnvC proteins and is located at what is expected to be a knob position in heptad XXII (see **Fig. 1b**).

**Figure 4.**
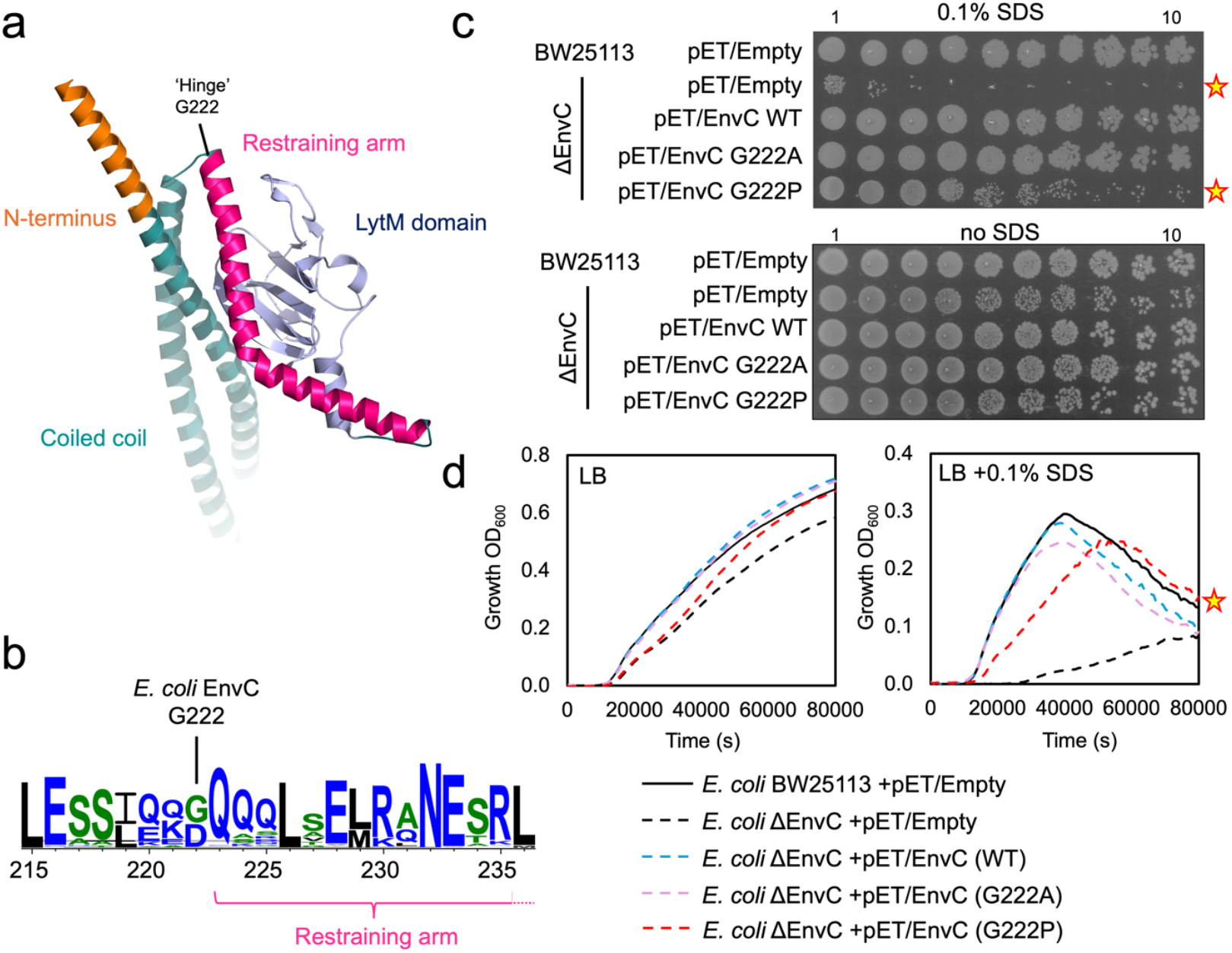
Mutations in the molecular hinge of EnvC. (a) Structural figure showing the EnvC hinge. (b) Weblogo representations of residue conservation at the EnvC hinge. (c) Detergent sensitivity assays for strains dependent on EnvC hinge variants. 10-fold serial dilutions of the indicated cultures are spotted from left-to-right on an LB agar plate containing 0.1% SDS. Stars indicate reduced detergent viability for strains that lack EnvC or are solely dependent on an EnvC Gly222Pro variant. (d) Growth curves for indicated strains grown in LB or LB supplemented with SDS detergent. All growth curves are the mean of at least 3 repeats.

The additional flexibility and the absence of a conventional ‘knob’-like sidechain led us to further investigate the importance of Gly222. We first made alanine and proline substitutions and tested their ability to complement the cell envelope defects of an EnvC-deficient strain. Alanine has a high propensity to form alpha helical secondary structures (29) thus we predicted the Gly222Ala variant may have a tendency towards activation. On the other hand, Proline, due to its sidechain ring structure, lacks the backbone N-H group needed to form an alpha helical structure, which might potentially block activation. *E. coli* cells lacking EnvC have reduced viability on SDS agar due to cell envelope defects. Complementation with either the wild type protein or Gly222Ala EnvC restored viability on SDS agar to the wild type levels, while the Gly222Pro variant only partly complemented EnvC deficiency (**Fig. 4c**). Similar results were found for growth in LB broth containing detergent (**Fig. 4d**) suggesting that EnvC Gly222 variants have a modestly impaired function compared to wild type.

While the effect of each Gly222 variant was broadly in line with our predictions, the magnitude of these phenotypic differences was small – especially in comparison to other mutations established in the FtsEX-EnvC-AmiA system. We therefore conclude that Gly222 may have some minor mechanistic importance as part of the EnvC hinge, but it is not an essential residue for the FtsEX-EnvC activation mechanism.

### The EnvC N-terminus is necessary for amidase activation

The N-terminus of EnvC consists of an exposed portion of alpha helix that contains three heptad repeats. In the autoinhibited conformation, these three heptads are unpaired and face the solvent (**Fig. 5a**), but in the activated conformation, these heptads have been predicted to pair with heptads in the restraining arm (8). The reversible interaction between the EnvC N-terminus and the restraining arm is a core prediction of the currently understood mechanism of conformational change that underpins the recruitment and activation of amidases by FtsEX-EnvC but, to our knowledge, this has not been tested by mutagenesis.

**Figure 5.**
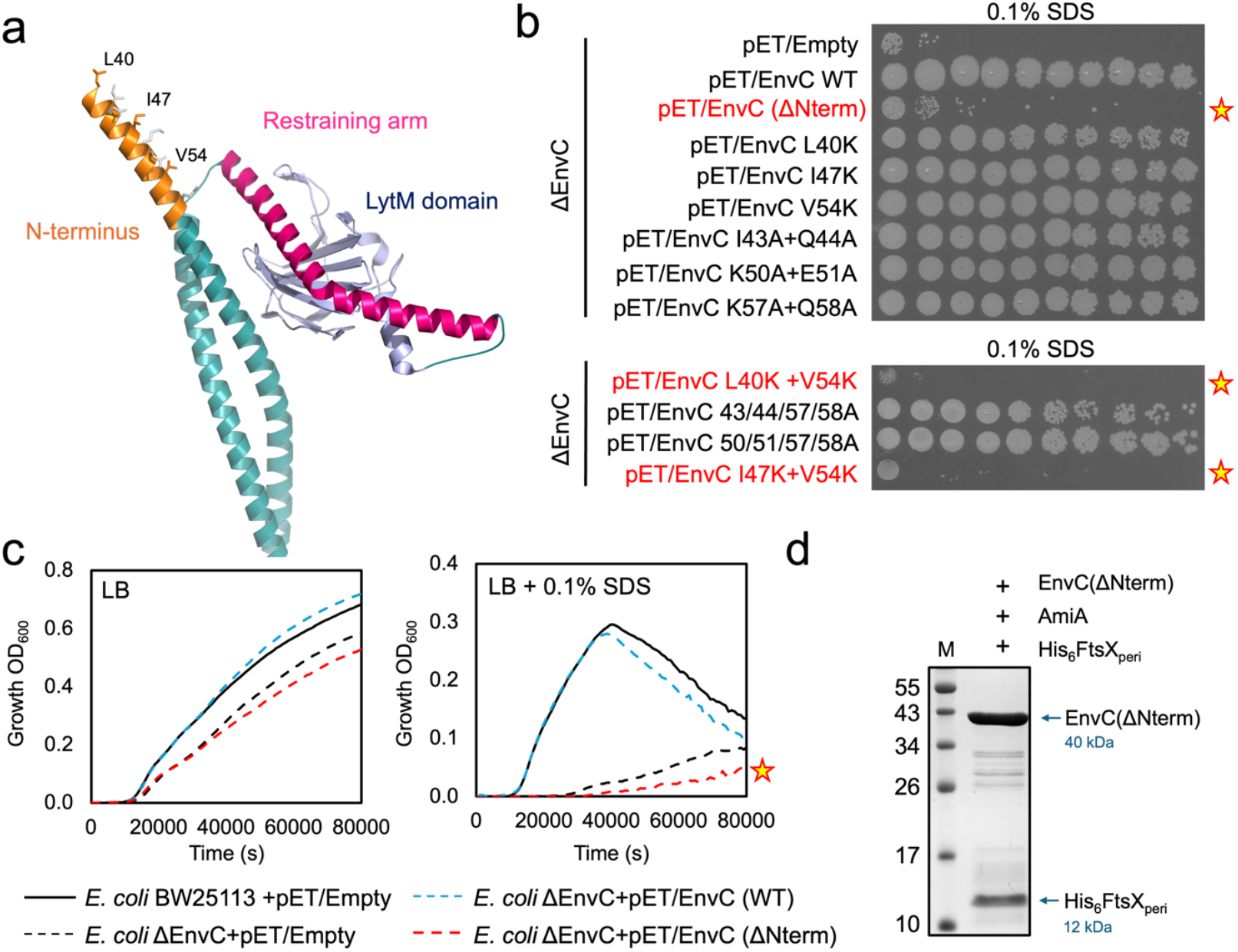
The EnvC N-terminus is required for activation of FtsEX-EnvC. (a) Close-up view of the EnvC N-terminus. (b) Detergent viability assays for strains dependent on plasmid-based expression of indicated EnvC variants. Stars indicate reduced viability on SDS agar due to mutations in the EnvC N-terminus. (c) Growth curves in LB broth containing SDS detergent. A star indicates an SDS sensitive strain that is dependent on EnvC lacking the N-terminus. (d) Purification of the EnvC(ΔNterm) variant in complex with the FtsX periplasmic domain in the presence of AmiA. All growth curves are the mean of multiple repeats.

To experimentally test the importance of the EnvC N-terminus, we made mutations in the N-terminal heptads and explored each variant’s ability to complement the SDS sensitivity phenotype of an *envc* knockout strain (**Fig. 5b**). Amino acid substitutions were chosen to disrupt the predicted knobs-into-holes packing expected to underpin the interaction of the N-terminus with the restraining arm. We also generated an EnvC construct that lacks the N-terminus completely – EnvC(ΔNterm). When expressed in the bacterial periplasm via pET22-based vector, the wild type EnvC protein completely rescues SDS sensitivity of the *envc* knockout strain. However, cells expressing EnvC without the N-terminus remain sensitive to SDS suggesting this feature is essential for EnvC function.

To further test the hypothesised importance of individual heptads, we assessed point mutations in the EnvC N-terminus (**Fig. 5b**). Individual disruption of the ‘knob’ residues from each of the three N-terminal heptads (L40K, I47K and V54K) did not oblate EnvC function and neither did individual disruption of any of the three residue pairs (I43A+Q44A, K50A+E51A and K57A+Q58A) that form the ‘holes’ within each heptad. However, two EnvC variants lacking pairs of knob residues (L40K+V54K and I47K+V54K) were unable to complement *envC* deficiency lending credence to the hypothesised knobs-into-holes packing that underpins activation of EnvC.

We also tested whether the EnvC N-terminal deletion variant could complement *envc* deficiency on the knockout strain in LB broth containing SDS (**Fig. 5c**). Expression of wild type EnvC completely complements the SDS sensitive phenotype while the protein lacking the N-terminus was unable to do so.

To control for the possibility that disruption of the N-terminus might simply impair the stability of EnvC, we made an EnvC expression construct in which the N-terminus was deleted and co-expressed it alongside the periplasmic domain of FtsX. The EnvC(ΔNterm)-FtsX periplasmic domain complex purifies well showing that deletion of the N-terminus does not disrupt the expression and folding of EnvC and that it remains capable of binding to FtsX (**Fig. 5d**). This construct also remains unable to bind AmiA, showing that removal of the N-terminus does not disrupt the EnvC autoinhibition mechanism (**Fig. 5d**). Taken together, these data support the hypothesised importance of the EnvC N-terminal heptad repeats in the activation mechanism of FtsEX-EnvC.

### Cysteine-locking of EnvC in its amidase-activating conformation

To further test the hypothesised knobs-into-holes packing expected in the activated FtsEX-EnvC complex, we attempted to trap the activated complex using a cysteine locking experiment. Based on modelling of the activated complex as a perfect coiled coil, we introduced pairs of cysteine residues into the EnvC N-terminus and restraining arm with the anticipation that these would form disulfide bonds when expressed in the periplasmic space. While we initially predicted that cysteine locking would be most effective between cognate ‘knob’ and ‘hole’ residues, our *in silico* modelling suggested that opposing ‘hole’ residues may in fact be better suited for optimal disulfide geometry (**Fig. 6a**). After some experimentation, we identified a cysteine pair variant (R37C+I240C) that appears to fit the bill. When expressed in the periplasm of a strain lacking the chromosomal copy of *envc*, the R37C+I240C EnvC variant renders cells sensitive to SDS even though the corresponding single cysteine variants (R37C, I240C) are unaffected (**Fig. 6b**). A second locking pair, E36C+I1240C, was also identified (**Fig. 6b**) although this was not analysed further.

**Figure 6.**
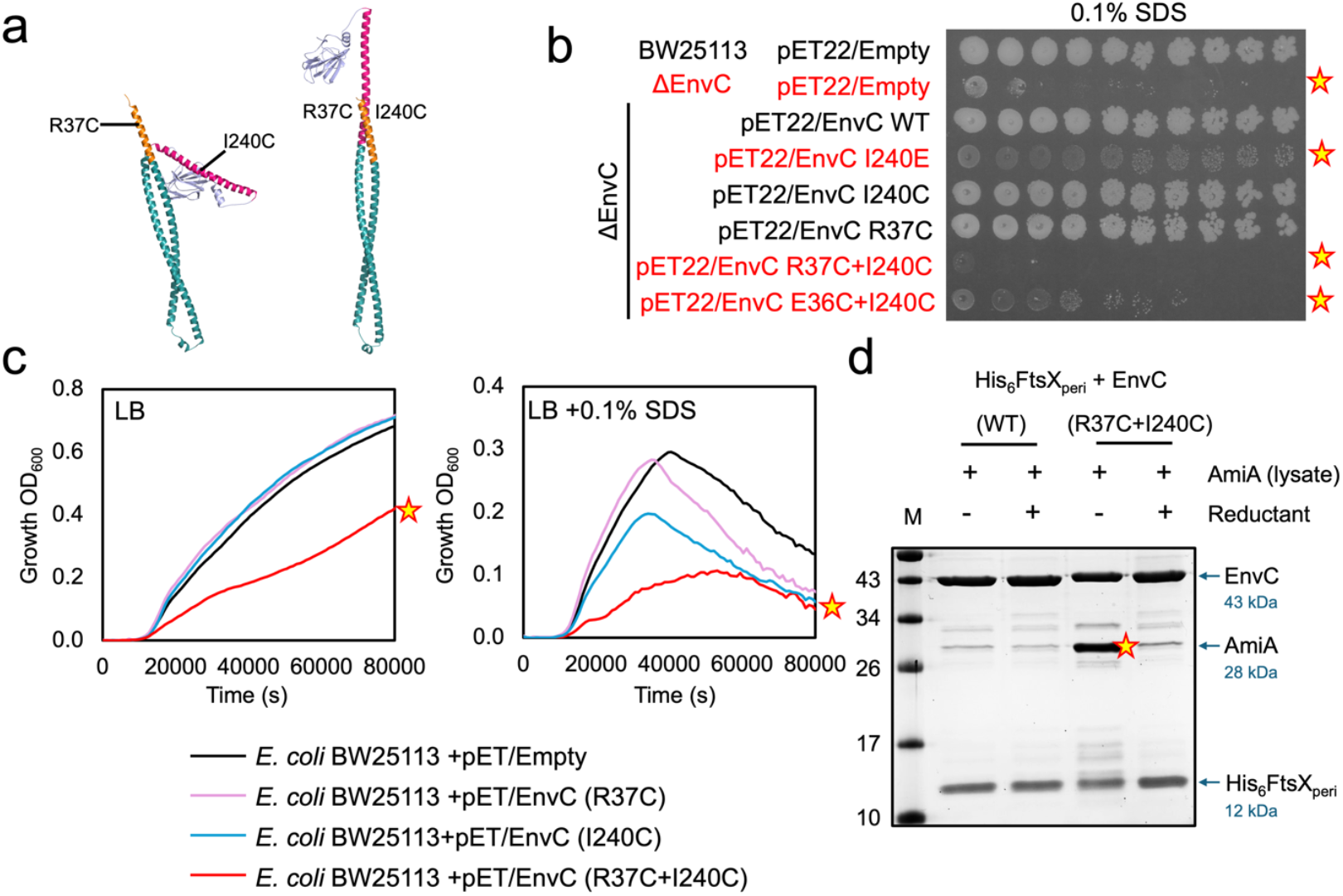
Disulfide locking of EnvC in an active conformation using engineered cysteine residues in the N-terminus and restraining arm. (a) Structural figure indicating the proposed crosslink position in EnvC. (b) Detergent sensitivity assay for strains grown on SDS agar. A 10-fold serial dilution for each culture was spotted on the agar in series from left to right starting at OD_600_ 1. Stars indicate detergent sensitive strains. (c) Growth curves in LB or LB broth containing SDS detergent. Star indicates growth of a strain expressing the EnvC(R37C+I240C) cross-linkable variant. (d) Co-purifications of the cross-linkable EnvC(R37C+I240C)-FtsX periplasmic domain complex in the presence of AmiA. Only the FtsX periplasmic domain is His-tagged. AmiA co-purifies with the cross-linked variant under oxidising conditions but not in the presence of reductant (TCEP).

Subsequent experiments in LB broth showed that expression of R37C+I240C EnvC not only fails to complement the cell envelope defect of the *envc* knockout, but also actively impairs the growth of wild type cells consistent with improper activation of amidases (**Fig. 6c**). In fact, a wild type *E. coli* strain expressing the R37C+I240C EnvC variant shows a greater growth defect in LB than a strain that lacks EnvC. These data suggest that R37C+I240C very likely forms a disulfide bond and can activate amidases in the periplasmic space.

It is noteworthy that the I240C mutation used for disulfide locking the restraining arm to R37C in the N-terminus is the same position as the I240E mutation used to disrupt the autoinhibition mechanism. This makes comparison of the I240E and I240C single mutants particularly insightful as they each have distinct properties. Complementation of the EnvC deficient strain with the I240C variant completely rescues growth to the level of the wild type EnvC while the I240E mutation exacerbates growth defects. Furthermore, expression of I240C in wild type cells has no impact on growth, while expression of I240E causes growth defects. When grown on LB agar containing SDS, the EnvC-deficient strain expressing I240E EnvC produces translucent colonies that are noticeably smaller than wild type while the I240C variant rescues the EnvC deficiency and appear completely normal (**Fig. 6b**). These comparisons demonstrate that I240C alone does not disrupt autoinhibition, while I240E does. The different behaviour of the two I240 mutations is explained by the nature of the substitutions - a cysteine residue is easily accommodated within the amidase-binding site, while the charged glutamate (I240E) is incompatible with the available space and hydrophobic character of the groove.

To further test the hypothesis that the cysteine-locked variant binds and activates amidases, we purified R37C+I240C EnvC in complex with the FtsX periplasmic domain and tested its ability to bind AmiA compared to the wild type in both the presence and absence of a disulfide-breaking reductant (**Fig. 6d**). We found that the soluble EnvC-FtsX periplasmic domain complex has minimal affinity for AmiA (consistent with autoinhibition) and is unaffected by the presence or absence of reductant (**Fig. 6d** *left*). However, the R37C+I240C variant binds AmiA strongly in the absence of reductant, but not at all when reductant is present (**Fig. 6d** *right*). Reversibility of the amidase binding function of the cysteine-locking variant due to the addition of a disulfide-breaking reductant indicates that the R37C+I240C variant only binds the amidase when a disulfide bond is present between the N-terminus and the restraining arm. The experiment also confirms that amidase binding cannot be explained by the I240C mutation dislodging the restraining arm as the R37C+I240C variant does not bind the amidase under reducing conditions. The combined data from *in vivo* and *in vitro* are consistent with R37C+I204C forming an intramolecular disulfide bond that locks the protein in an activated conformation allowing amidase recruitment and activation.

## Discussion

FtsEX-EnvC has a well-understood molecular mechanism of amidase activation built on many years of research from numerous groups (**Figure 1**). Here we have experimentally tested key aspects of the proposed molecular mechanism using site-directed mutagenesis. We show that the EnvC autoinhibition mechanism relies on hydrophobic interactions between the restraining arm and the amidase-binding groove of the LytM domain that can be disrupted by single amino acid mutations (**Figures 2**). These mutations overcome the EnvC autoinhibition mechanism and facilitate inauthentic recruitment of both AmiA and AmiB. Residue Ile240 appears especially important with charged residue substitutions producing growth defects when expressed in *E. coli* (**Figure 3**). We also probed the role of Gly222 within the EnvC ‘hinge’ that is expected to underpin the conformational change between autoinhibited and amidase-recruiting conformations (**Figure 4**). Proline substitution of Gly222 lightly hinders EnvC function, but the glycine does not appear to be essential. We also challenged the hypothesised knobs-into-holes packing that is suspected to occur between the restraining arm and the protein N-terminus during activation of EnvC (**Figure 5**). Deletion of the N-terminus, or disruption of the heptad repeats within it, blocks EnvC’s amidase activation function. Finally, we show that introducing a disulfide bond between the restraining arm and the EnvC N-terminus renders EnvC prone to continuous activation facilitating the unimpeded binding of amidases (**Figure 6**). Collectively, these experiments provide ample support for the currently understood mechanism of amidase activation by FtsEX-EnvC and underline the importance of both the EnvC N-terminus and restraining arm for regulation of periplasmic peptidoglycan amidases.

The mechanism by which FtsEX-EnvC couples ATP binding and hydrolysis to the activation of periplasmic amidases is now understood in extensive molecular detail and supported by numerous experiments and structural observations. However, there remain aspects of the wider FtsEX-EnvC mechanism that need to be further explored. Most pressingly, the current understanding of amidase regulation is that amidases are controlled by a series of nested autoinhibition mechanisms that are ultimately relieved by the binding and hydrolysis of ATP by FtsEX, however, it remains unclear what, if anything, regulates ATP binding and hydrolysis on the cytoplasmic side of the membrane. Similarly, while it is understood that FtsEX-EnvC is recruited to the division site by interactions with the Z-ring (11), the molecular nature of these interactions remains to be deduced. Future experiments are needed to establish precisely how FtsEX-EnvC interacts with other components of the cell division machinery and how these interactions regulate the ATP binding and hydrolysis cycle underpinning amidase activation in the periplasm.

Looking beyond *E. coli*, there is plentiful variety in the mechanisms of FtsEX-EnvC -like systems identified across the bacterial kingdom. This is especially true of FtsEX-EnvC -like systems such as FtsEX-RipC and FtsEX-PcsB where the EnvC-like components are enzymatically active and therefore do not recruit a separate amidase to begin peptidoglycan hydrolysis (12,26,30–32). Recent structures of the *Mycobacterium tuberculosis* FtsEX-RipC system suggests that RipC lies flat in the inactive conformation and is straightened upon activation by ATP-driven FtsEX mechanotransmission (26). It remains unclear whether these systems use autoinhibition mechanisms to regulate their enzymatic domains or whether it is solely the localisation of their enzymatic domains relative to the peptidoglycan layer that determines their processivity.

One practical use for building a better understanding of FtsEX-EnvC-amidase system is to develop inhibitors that block cell division or cause uncontrolled activation of the peptidoglycan amidases. There are now several mutations in the FtsEX-EnvC-AmiA/B system that emulate the effect of such hypothetical molecules. For example, mutations in the ATP-binding site that interfere with nucleotide binding (K41A) or hydrolysis (E163Q), or mutations that weaken the interaction between FtsEX and EnvC (eg F152E in FtsX), each block the amidase activation mechanism and have phenotypes that mimic cells lacking the genes encoding FtsE, FtsX or EnvC (8). Deletion or disruption of the EnvC N-terminus has a similar effect, preventing the activation of amidases in a genetic background that lacks the wild type EnvC. Conversely, mutations in EnvC that relieve the autoinhibition mechanism (such as the EnvC I240E mutation and disulfide-locking R37C+I240C mutations reported here) promote promiscuous activation of peptidoglycan amidases causing growth and cell envelope defects - even when expressed in the wild type genetic background. Other (dissimilar) ‘dominant’ variants have been reported previously - for example, the expression of amidases lacking their internal autoinhibition helix (or bearing point mutations that loosen its grip) can cause bacterial lysis (9,18,19). However, amidase variants and both the I240E and R37C+I240C variants reported here are not as effective in producing defects as the isolated EnvC LytM domain (8,22) (which can freely bind and activate multiple amidases in the periplasm without regulation by FtsEX) perhaps suggesting there are additional layers of amidase regulation that are not yet fully appreciated. Notably however, none of these variants seem to render the bacterium completely inviable suggesting that cells possess robust compensatory mechanisms that can, at least partially, mitigate the effect of promiscuous amidase activation. It therefore remains unclear whether targeting FtsEX-EnvC or the amidases represents a viable strategy for drug development.

In summary, we investigated key structural features of EnvC that are proposed to underpin an ATP-driven conformational change in the FtsEX-EnvC complex that regulates the binding and activation of periplasmic peptidoglycan amidases during cell division. We found extensive support for the reversible knobs-into-holes conformational change in EnvC and identified both the protein N-terminus and Ile240 within the restraining arm as key mechanistic residues involved in both autoinhibition and amidase activation by FtsEX-EnvC.

## Methods

### DNA constructs

A table describing the DNA constructs used in this report appears in the Supplemental Data (**Table S1**). For cloning, PCR typically used Q5 based DNA polymerase and primers engineered to include restriction sites at their 5’ end. Cloning then used standard restriction enzymes for cutting DNA and T4 DNA ligase for assembly. EnvC constructs lacking their N-terminus were generated by PCR starting at base 111 (coding for Arg55) and ligated into an appropriate pET vector (eg pET22 for periplasmic expression, or pET-Duet for cytoplasmic co-expression with the FtsX periplasmic domain). Point mutations were introduced using Quikchange Site-directed Mutagenesis and sub-cloned into target vectors using restriction-based cloning methods. All DNA constructs were confirmed by DNA sequencing (Genewiz).

### Bacterial 2-hybrid assays

Protein-protein interactions were monitored using the BACTH system (33). Assays were performed in *E. coli* BTH101 cells transformed with compatible plasmid pairs. After transformation, cell cultures were grown overnight at 30 °C in LB supplemented with 50 µg/mL ampicillin and 25 µg/mL kanamycin. Cultures were then spotted onto LB agar plates containing 40 µg/mL X-gal, 0.5 mM IPTG, 50 µg/mL ampicillin and 25 µg/mL kanamycin. Agar plates were incubated at 20 °C for 3 days and imaged using an Epson document scanner.

### Protein co-purification assays

*E. coli* C43(DE3) transformed with pET-based plasmids encoding AmiA or EnvC variants with/without a periplasmic domain of FtsX were grown at 30 °C in 2YT media. Protein expression was induced at OD 0.6 using 1 mM IPTG and progressed overnight at 30 °C. Cells were pelleted by centrifugation (10 min, 6,000 ×*g*), resuspended in Buffer A (26 mM Imidazole, 300 mM NaCl, 50 mM HEPES pH 7.2) and then broken by sonication (seven 30 s pulses using a Bandelin Sonopuls instrument). Lysates were clarified by centrifugation (20 min, 30,000 ×*g*) and mixed as required for each experiment. Mixed lysates were then incubated with Bio-Rad Ni-IMAC resin overnight at 4 °C with continuous agitation via a tube roller. Each protein-bound IMAC resin was transferred to an empty PD10 column, and extensively washed with the same buffer used for resuspension. Remaining proteins were then eluted from the resin with a high imidazole buffer (Buffer B: 250 mM Imidazole, 300 mM NaCl 50 mM HEPES pH 7.5) and analysed by SDS-PAGE.

For cysteine locking experiments, purifications were performed similarly with modifications to account for the use of TCEP. Briefly, the EnvC-FtsX periplasmic domain complex (and equivalent R37C+I240C variant) were each prepared as a cleared cell lysate and split into two equal volumes. 5 mM TCEP was added to one fraction and omitted from the other. After 1 hr agitation at 4 °C, both volumes were mixed with a cleared lysate containing untagged AmiA (with or without TCEP). Each volume was then incubated with Bio-Rad Ni-IMAC resin overnight at 4 °C. Bound proteins were washed with 20 column volumes of wash Buffer (35 mM Imidazole, 300 mM NaCl, 50 mM HEPES pH 7.2) supplemented with 5mM TCEP as required). A final wash was performed using wash buffer without TCEP before eluting with Buffer B. Eluted proteins were separated using SDS-PAGE.

### EnvC complementation experiments

Complementation experiments used *E. coli* BW25113 or an otherwise isogenic *envc* knockout strain from the Keio collection. BW25113 and *Δenvc* strains were transformed by electroporation with pET22-based vectors designed to express either wild type or variant EnvC proteins in the periplasm. An empty pET22 plasmid was used as a negative control. Cell envelope integrity of each strain was assessed using sensitivity to SDS detergent on both solid agar and in LB broth. Briefly, cells were grown in LB supplemented with 50 µg/mL ampicillin and adjusted to an OD of 1 (at 600 nm) before serial dilution in fresh LB (10-fold each time). Each dilution series was then spotted onto solid LB agar plates (1 mM IPTG and 50 µg/mL ampicillin), made with, or without, 0.1 % SDS and grown overnight at 37 °C before imaging via an Epson document scanner. For liquid broth experiments, OD 1 cultures were diluted 1000-fold and a 20 µL volume used to seed 200 µL cultures of LB with 50 µg/mL ampicillin and 1 mM IPTG (or additionally supplemented with 0.1 % SDS) in 96-well plates. Growth curves were then measured at 37 °C using the OD at 600 nm as a proxy for bacterial proliferation. Measurements were taken every 15 minutes using a shaking plate reader (Thermo Multiskan Sky). All growth curves are the average of at least 3 repeats.

### Structure analysis and bioinformatics

Homologous sequences for EnvC were identified using the Kegg database (34). Multiple sequence alignments were constructed with Kalign (35) and visualised using Weblogo (36). Structural figures were generated using Pymol (37).

## Acknowledgements

We thank the Warwick Media Kitchen for making bacterial culture media and the School of Life Sciences Tech Team for laboratory support. The work was funded by the BBSRC (BB/V017101/1) with support from the Howard Dalton Centre.

## Supplemental Information

**Table S1.** Plasmids used in this study.

| Plasmid | Vector | Contents | Label in Figure | Figure |
| --- | --- | --- | --- | --- |
| pJC6 192 | pUT18C | FtsX (110-209) -T18<br>*FtsX periplasmic domain for B2H* | FtsX<sub>peri</sub>-T18 | 2b,2c |
| pTB1 011 | pUT18C | AmiA (35-289) -T18<br>*AmiA for B2H* | AmiA-T18 | 2b,2c |
| pJC6 238 | pUT18C | AmiB (23-445) -T18<br>*AmiB for B2H* | AmiB-T18 | 2b |
| pJC6 124 | pKNT25 | EnvC (35-419) -T25<br>*EnvC for B2H* | EnvC-T25 | 2b,2c |
| pJC6 257 | pKNT25 | EnvC (278-419) -T25<br>*EnvC lytM domain for B2H* | EnvC<sub>lytM</sub>-T25 | 2b,2c |
| pJC6 743 | pKNT25 | EnvC (35-419) -T25 (L236E) | EnvC-T25 L236E | 2b |
| pJC6 745 | pKNT25 | EnvC (35-419) -T25 (L236K) | EnvC-T25 L236K | 2b |
| pJC6 747 | pKNT25 | EnvC (35-419) -T25 (I240E) | EnvC-T25 I240E | 2b |
| pJC6 751 | pKNT25 | EnvC (35-419) -T25 (I240K) | EnvC-T25 I240K | 2b |
| pJC6 753 | pKNT25 | EnvC (35-419) -T25 (E233R) | EnvC-T25 E233R | 2c |
| pJC6 757 | pKNT25 | EnvC (35-419) -T25 (R237E) | EnvC-T25 R237E | 2c |
| pJC6 761 | pKNT25 | EnvC (35-419) -T25 (E233R+R237E) | EnvC-T25 233+237 | 2c |
| - | pUT18C | Empty vector control for B2H<br>T18 fragment | T18 | 2b,2c |
| - | pKNT25 | Empty vector control for B2H<br>T25 fragment | T25 | 2b,2c |
| - | pET22 | Empty vector control for complementation<br>experiments | pET/Empty | 3a,3b,4c,4d |
| pJC6 260 | pET22 | EnvC (35-419)<br>*EnvC with pelB signal for periplasmic expression* | pET/EnvC (WT) | 3a,3b,4c,4d,5b,5c,6b |
| pJC6 957 | pET22 | EnvC (35-419) (I240E)<br>*EnvC with pelB signal for periplasmic expression* | pET/EnvC (I240E) | 3a,3b,4c,6b |
| pJC6 960 | pET22 | EnvC (55-419)<br>*EnvC without mature N-terminus, with pelB signal for periplasmic expression* | pET/EnvC (ΔNterm) | 5b,5c |
| pJC6 986 | pET22 | EnvC (35-419) (R37C)<br>*with pelB signal for periplasmic expression* | pET/EnvC R37C | 6b,6c |
| pJC6 994 | pET22 | EnvC (35-419) (I240C)<br>*with pelB signal for periplasmic expression* | pET/EnvC I240C | 6b,6c |
| pJC6 987 | pET22 | EnvC (35-419) (R37C+I240C)<br>*with pelB signal for periplasmic expression* | pET/EnvC R37C+I240C | 6b,6c |
| pJC6 988 | pET22 | EnvC (35-419) (E36C+I240C)<br>*with pelB signal for periplasmic expression* | pET/EnvC E36C+I240C | 6b |
| pJC7 166 | pET22 | EnvC (35-419) (G222A)<br>*Hinge mutant for periplasmic expression* | pET/EnvC G222A | 4c,4d |
| pJC7 202 | pET22 | EnvC (35-419) (L40K) | pET/EnvC L40K | 5b |
| pJC7 204 | pET22 | EnvC (35-419) (I43A+Q44A) | pET/EnvC I43A+Q44A | 5b |
| pJC7 234 | pET22 | EnvC (35-419) (I47K) | pET/EnvC I47K | 5b |
| pJC7 236 | pET22 | EnvC (35-419) (V54K) | pET/EnvC V54K | 5b |
| pJC7 238 | pET22 | EnvC (35-419) (K57A+Q58A) | pET/EnvC K57A+Q58A | 5b |
| pJC7 244 | pET22 | EnvC (35-419) (K50A+E51A) | pET/EnvC K50A+E51A | 5b |
| pJC7 275 | pET22 | EnvC (35-419) (L40K+V54K) | pET/EnvC L40K+V54K | 5b |
| pJC7 277 | pET22 | EnvC (35-419) (I43A+Q44A+K57A+Q58A) | pET/EnvC 43/44/57/58 | 5b |
| pJC7 285 | pET22 | EnvC (35-419) (K50A+E51A+K57A+Q58A) | pET/EnvC 50/51/57/58 | 5b |
| pJC7 291 | pET22 | EnvC (35-419) (I47K+V54K) | pET/EnvC I47K+V54K | 5b |
| pJC7 322 | pET22 | EnvC (35-419) (G222P)<br>*Hinge mutant for periplasmic expression* | pET/EnvC G222P | 4c,4d |
| pJC6 763 | pET21 | AmiA (35-289)<br>*No His-tag, for in vitro protein binding experiments* | AmiA | 2d,2e,5d,6d |
| pJC6 474 | pETDuet1 | EnvC (222-419)<br>*N-terminal His tag, no secretion signal, no coiled coil domain, for in vitro protein experiments* | His<sub>6</sub>-EnvC(ΔCC) WT | 2e |
| pJC6 775 | pETDuet1 | EnvC (222-419) (I240E)<br>*N-terminal His-tag, no secretion signal, no coiled coil domain, for in vitro protein experiments* | His<sub>6</sub>-EnvC(ΔCC) I240E | 2e |
| pJC6 120 | pETDuet1 | FtsX (110-209) + EnvC (35-419)<br>*Co-expression system for the EnvC-FtsX periplasmic domain complex. His-tag on c-terminus of FtsX periplasmic domain. For in vitro protein experiments* | His<sub>6</sub>-FtsXperi and EnvC | 2d,6d |
| pJC6 766 | pETDuet1 | FtsX peri (110-209) + EnvC (35-419) I240E<br>*Co-expression system for the EnvC-FtsX periplasmic domain complex. His-tag on c-terminus of FtsX periplasmic domain. For in vitro protein experiments* | His<sub>6</sub>-FtsXperi and EnvC I240E | 2d |
| pJC6 968 | pETDuet1 | FtsX peri (110-209) + EnvC (R37C+I240C)<br>*Co-expression system for the EnvC-FtsX periplasmic domain complex. His-tag on c-terminus of FtsX periplasmic domain. For in vitro protein experiments* | His<sub>6</sub>-FtsXperi and EnvC (R37C+I240C) | 6d |
| pJC7 148 | pETDuet1 | FtsX peri (110-209) + EnvC (55-419)<br>*Co-expression system for the EnvC(ΔNterm) -FtsX periplasmic domain complex. EnvC is missing the mature N-terminus. His-tag on c-terminus of FtsX periplasmic domain. For in vitro protein experiments* | His<sub>6</sub>-FtsXperi and EnvC (ΔNterm) | 5d |

